# Subfamily-specific differential contribution of individual monomers and the tether sequence to mouse L1 promoter activity

**DOI:** 10.1101/2021.12.03.471143

**Authors:** Lingqi Kong, Karabi Saha, Yuchi Hu, Jada N. Tschetter, Chase E. Habben, Leanne S. Whitmore, Changfeng Yao, Xijin Ge, Ping Ye, Simon J. Newkirk, Wenfeng An

## Abstract

**Background:** The internal promoter in L1 5’UTR is critical for autonomous L1 transcription and initiating retrotransposition. Unlike the human genome, which features one contemporarily active subfamily, four subfamilies (A_I, Gf_I and Tf_I/II) have been amplifying in the mouse genome in the last one million years. Moreover, mouse L1 5’UTRs are organized into tandem repeats called monomers, which are separated from ORF1 by a tether domain. In this study, we aim to compare promoter activities across young mouse L1 subfamilies and investigate the contribution of individual monomers and the tether sequence.

**Results:** We observed an inverse relationship between subfamily age and the average number of monomers among evolutionarily young mouse L1 subfamilies. The youngest subgroup (A_I and Tf_I/II) on average carry 3-4 monomers in the 5’UTR. Using a single-vector dual-luciferase reporter assay, we compared promoter activities across six L1 subfamilies (A_I/II, Gf_I and Tf_I/II/III) and established their antisense promoter activities in a mouse embryonic fibroblast cell line. Using consensus promoter sequences for three subfamilies (A_I, Gf_I and Tf_I), we dissected the differential roles of individual monomers and the tether domain in L1 promoter activity. We validated that, across multiple subfamilies, the second monomer consistently enhances the overall promoter activity. For individual promoter components, monomer 2 is consistently more active than the corresponding monomer 1 and/or the tether for each subfamily. Importantly, we revealed intricate interactions between monomer 2, monomer 1 and tether domains in a subfamily-specific manner. Furthermore, using three-monomer 5’UTRs, we established a complex nonlinear relationship between the length of the outmost monomer and the overall promoter activity.

**Conclusions:** The laboratory mouse is an important mammalian model system for human diseases as well as L1 biology. Our study extends previous findings and represents an important step toward a better understanding of the molecular mechanism controlling mouse L1 transcription as well as L1’s impact on development and disease.

## Introduction

Long interspersed elements type 1 (LINE1s, or L1) are ubiquitous non-long terminal repeat (LTR) retrotransposons in mammals (1, 2), comprising 17% and 19% of the human and mouse genome, respectively (3, 4). Only a very small fraction of genomic L1 copies are full-length as the vast majority of L1s suffer “structural defects”, such as 5’-truncation (5, 6), 5’-inversion (6-8), or internal rearrangement (9). A full-length L1 is 6-7 kb long (10, 11), encompassing a 5’ untranslated region (5’UTR), two open reading frames (ORF1 and ORF2) and a 3’ untranslated region (3’UTR). The 5’UTR contains an internal promoter, which is critical for autonomous L1 transcription (12-14) and the initiation of L1 retrotransposition. The resulting L1 mRNA serves dual functions. First, it can be translated into two L1 proteins (ORF1p and ORF2p); both are essential for L1 retrotransposition (15, 16). Second, the same L1 mRNA is the preferred template for ORF2p-mediated reverse transcription over other cellular RNAs, in a phenomenon known as cis preference (17, 18). Based on comprehensive surveys of full-length elements among recently integrated human L1s (19, 20), approximately 30% of the new L1 insertions are full-length loci, which can potentially prime additional rounds of retrotransposition from their 5’UTRs.

Genomic L1 sequences are grouped into subfamilies according to their evolutionary history. Among L1s in the human genome, the oldest subfamilies L1MA to L1ME are shared with other mammals, but the younger L1PB and L1PA subfamilies are only found in primates. The youngest subfamily, L1PA1 (also called L1Hs), is specific to humans (21). A remarkable feature of L1 evolution is that new subfamilies frequently emerged by acquiring distinct 5’UTRs unrelated to those found in existing subfamilies (22). In the last ∼70 million years during primate evolution, there were at least eight episodes of 5’UTR replacement. It is believed that new 5’UTRs provide a mechanism for emergent subfamilies to avoid competition of host factors or to escape host suppression (22). The latest 5’UTR acquisition occurred ∼40 million years ago (MYA) in ancestral anthropoid primates and gave rise to subfamily L1PA8 (23). The overall architecture of this new 5’UTR had been maintained as a single lineage in later subfamilies from L1PA7 to L1PA1. Nevertheless, these subfamilies were subjected to continued host-L1 conflicts. For example, subfamilies L1PA6 to L1PA3 had evolved a ZNF93 binding motif in their 5’UTRs, which recruits ZNF93, triggering KAP1-mediated transcriptional silencing (24, 25). In contrast, a 129-bp deletion in the 5’UTR (inclusive of the binding site) allowed a subset of L1PA3, L1PA2, and L1PA1 to escape ZNF93 suppression (25). In addition, a single nucleotide change at position 333 created a functional m6A site, which first appeared in a subset of L1PA3 and then dominated in L1PA2 and L1PA1 (26). Primate L1 5’UTRs also possess an antisense promoter, which drives the expression of a third open reading frame (ORF0) as well as chimeric fusion transcripts with upstream cellular genes (27-29).

The laboratory mouse is an important mammalian model system for human diseases as well as L1 biology (30-33). Despite sharing many ancestral L1 subfamilies with the human genome, the mouse genome is dominated by lineage specific L1 subfamilies, which were initially evolved from ancestral L1MA6 elements ∼75 MYA at the divergence of the two species (4). A comprehensive analysis of full-length L1 sequences in the mouse genome identified 29 L1 subfamilies that have undergone amplification since the split between mouse and rat about 13 MYA (34). Overall, the evolution of mouse L1 subfamilies fits in the single lineage model as seen in the human genome. Similarly, young mouse L1 subfamilies frequently evolved by acquiring new 5’UTR sequences. Since the split from the rat, the mouse genome has experienced at least 11 episodes of 5’UTR replacement (34). The 29 L1 subfamilies feature seven types of 5’UTR sequences: Lx, V, Fanc, Mus, F, A and N (ordered by their first appearance in the genome from old to new) (34). The F type 5’UTR was resurrected from Fanc ∼6.4 MYA and led to the formation of subfamilies F_V to F_I, the youngest of which ceased amplification about 2 MYA. The A type 5’UTR was recruited approximately 4.6 MYA and appeared in seven L1 subfamilies (A_VII to A_I), with A_I being the youngest and active since 0.25 MYA. Remarkably, the F type 5’UTR had been revived three times through recombination of the 5’ portion of an F element with the 3’ portion of an A_III element, forming subfamilies Gf_II, Gf_I, and Tf_III/II/I respectively. As in the human genome, the evolutionary timeline of mouse L1s is also interspersed with episodes of multiple subfamilies coexisting over extended periods of time. For both human and mouse L1s, concurrently active subfamilies often possessed distinct 5’UTR promoter sequences (23, 34). This observation has led to a hypothesis that different promoters enabled subfamilies not to compete for the same transcription factors. Unlike the human genome, which features one contemporarily active subfamily, at least three subfamilies (Gf_I, Tf_I/II, and A_I) have been amplifying in the mouse genome in the last one million years (34, 35). Interestingly, phylogenetic evidence suggests that Gf_I and Tf_I/II in the laboratory mouse genome might be acquired through inter-specific hybridization rather than evolved from within its own genome (34). In any case, it is unclear whether all three subfamilies remain currently active in the germ line of the laboratory mice.

Owing to their lineage-specific nature human and mouse L1 5’UTRs share no sequence homology. Moreover, mouse L1 5’UTRs are distinctly different from human L1’s in that the former are organized into tandem repeats called monomers (11, 36). Such monomeric structures are also present in some other vertebrate L1s, including rat, hyrax, horse, elephant and opossum, but mouse L1 5’UTRs boast the highest number of monomers among all vertebrates (37). The number of monomers varies among individual L1s. For example, two recent full-length Tf insertions carried 5.7 and 7.5 monomers, respectively (38). Using reporter assays, it has been demonstrated 2-monomer is the minimal promoter structure to have significant transcriptional activity for L1spa, a Tf subfamily member (39). Similar tests have not been conducted for other mouse L1 subfamilies. Between monomers and ORF1 is a non-monomeric sequence, termed tether (40). In both A and Tf subfamily mouse promoters, tethers lacked significant transcriptional activity in reporter assays (39, 41). In this study, we aim to compare promoter activities across young mouse L1 subfamilies and investigate the contribution of individual monomers and the tether sequence using reporter assays.

## Results

### Most full-length L1s from young mouse L1 subfamilies possess two or more monomers

To profile mouse L1 promoter activities, we first analyzed the length distribution of mouse L1 5’UTRs by counting the number of monomers for full- or near full-length elements. Since elements from the old subfamilies would have accumulated numerous debilitating mutations, we limited our analysis to seven recently active subfamilies, including A_I, Tf_I, Tf_II, Gf_I, Tf_III, A_II, and A_III (listed from young to old). The estimated age for these L1 subfamilies ranges from 0.21 MYA for A_I to 2.15 MYA for A_III (Fig. 1A) (34). To tabulate elements carrying a specific number of monomers, L1 loci containing at least a partial 5’UTR are binned according to their respective 5’ start point (Fig. 1B). For example, if the 5’UTR of an element starts within the third monomer, it would be placed into the monomer 3 (M3) bin. We observed a trend of 5’UTR length shortening as subfamilies age. The vast majority of A_I elements (1032 out of 1125 or 91.7%), the youngest among this group, have at least two intact monomers. The distribution of A_I elements peaks at M3 (357 out of 1125 or 31.7%). In other words, more loci start within the third monomer than any other 5’UTR positions. In contrast, 87.6% (816/931) of the A_III loci, the oldest among this group, have fewer than two intact monomers, and 71.6% (667/931) of the loci start in monomer 2 (M2). This shortening trend is also evident if a comparison is made among closely related subfamilies (e.g., comparing among A_I, A_II and A_III, or among TF_I, Tf_II and Tf_III). Overall, among the loci with at least a partial 5’UTR from these seven mouse L1 subfamilies, 61.0% (3515/5765) have >2 intact monomers, 29.7% (1710/5765) have >3 intact monomers, 14.9% (858/5765) have >4 intact monomers, and 7.8% (230/5765) have >5 intact monomers. At the extreme end of the spectrum, there are seven loci that have >10 intact monomers (i.e., falling into M11+ bin), all belonging to A_I, Tf_I, Tf_II, and Gf_I subfamilies. To calculate the average number of monomers for each subfamily, we excluded loci with either >10 monomers or truncated within the tether (T) (Fig.1C). On average, L1 loci from the youngest subgroup carry >3 monomers (3.7, 3.5 and 3.1 monomers for A_I, Tf_I and Tf_II, respectively), followed by Gf_I (2.3 monomers), Tf_III (2.5 monomers), A_II (2.3 monomers), and A_III (1.5 monomers). An inverse relationship was observed between subfamily age and the average number of monomers among these seven mouse L1 subfamilies (simple linear regression: R = −0.91, p = 0.004).

**Figure 1.**
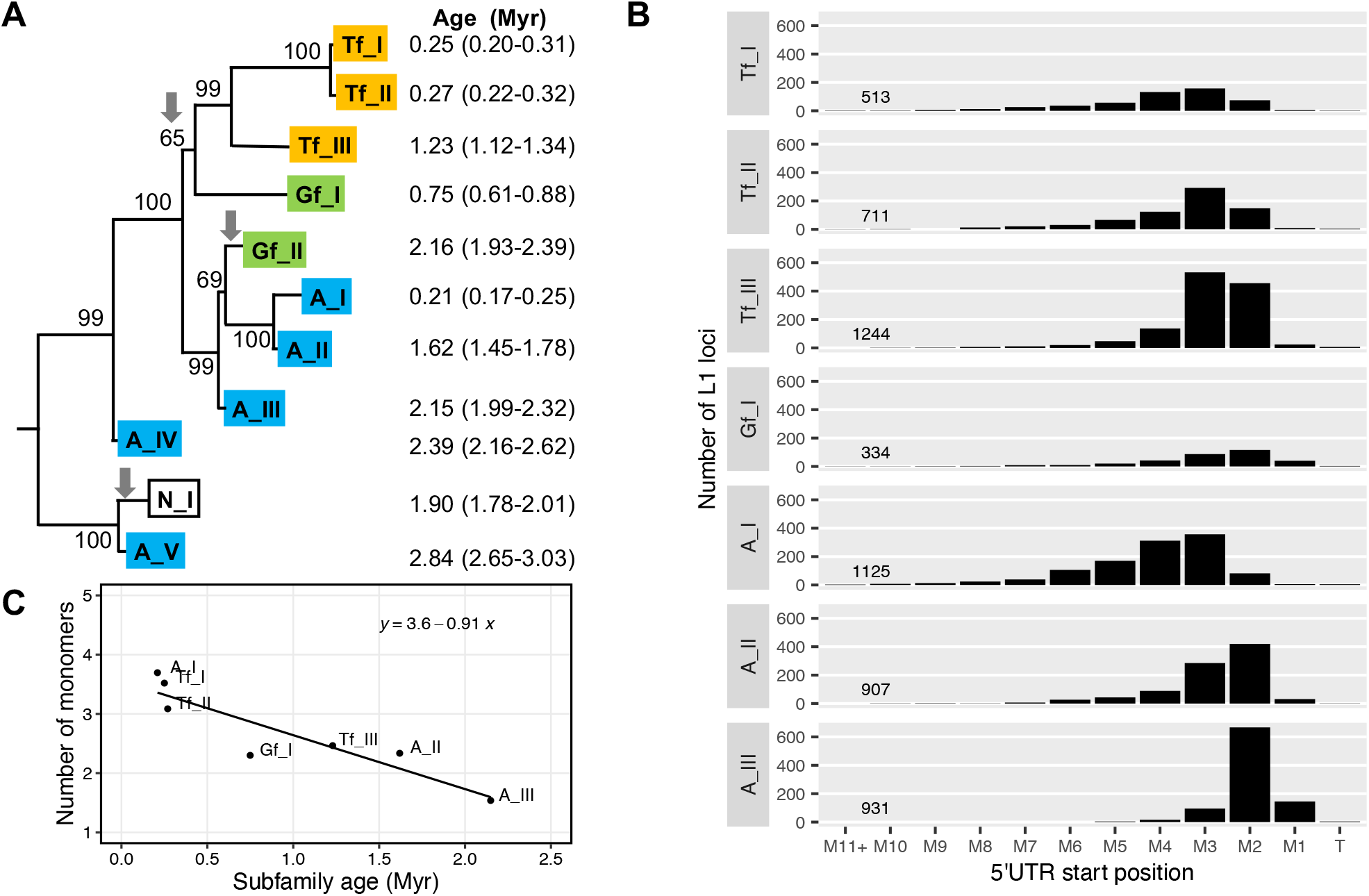
Phylogenetic relationship and promoter length distribution of young mouse L1 subfamilies. (**A**) A partial mouse L1 phylogenetic tree that consists of the youngest subfamilies. Adapted from Figure 1 of Sookdeo et al (34) under Creative Commons Attribution 4.0 International License (https://creativecommons.org/licenses/by/4.0/). The tree was built with the longest non-recombining region of ORF2 sequences using the maximum-likelihood method. The numbers indicate the percentage of time the labeled note was present in 1000 bootstrap replicates of the data. Downward arrows indicate the acquisition of a new 5’UTR. The age of each subfamily, in million years (Myr), was estimated by calculating the average pairwise divergence of the 3’UTR and converting the divergence to time assuming a neutral rodent genomic substitution rate of 1.1% per million year (see Table 1 of the original publication). We applied styling changes to highlight the Tf, Gf, and A subfamilies. (**B**) Distribution of the 5’UTR start position in different L1 subfamilies. For each subfamily, the number of L1 loci is tallied according to their starting nucleotide position relative to the tether (T), the first ten individual monomers (M1 to M10), M11 and beyond (M11+). (**C**) Inverse relationship between the average number of monomers and subfamily age. A simple linear regression line and the corresponding equation were shown along with individual data points.

### Two-monomer consensus sequences from six L1 subfamilies differ in their sense promoter activities

To quantitatively evaluate L1 promoter activity, we developed a single-vector dual-luciferase reporter assay (Fig.2A). In this vector design, a variant of L1 promoter drives the expression of firefly luciferase (Fluc), and an invariable HSV-TK promoter drives the expression of the Renilla luciferase (Rluc). The Rluc reporter cassette is embedded on the plasmid backbone as an internal control to normalize transfection efficiency. The L1 promoter activity is reported as the average Fluc/Rluc ratio among four replicate wells of NIH/3T3 cells. For this assay to work properly, it is important that Fluc and Rluc signals are both within the linear dynamic range (i.e., not saturated). Furthermore, there should be minimal crosstalk between the two reporter cassettes. To this end, we performed a titration experiment using varying amount of pCH117 plasmid per reaction in a 96-well assay format. Note the L1 promoter in the pCH117 plasmid was derived from an active human L1, L1RP (42). The Fluc and Rluc signals scaled proportionally to the amount of plasmid from 5 ng to 20 ng but started to plateau when 25 ng or more plasmid was used (Fig.2B). The Fluc/Rluc ratio was relatively stable within this range (Fig.2C). In subsequent assays, 10 ng plasmid DNA was used per well for all promoter assays.

To compare promoter activities across mouse L1 subfamilies, we first synthesized the consensus 5’UTR sequence of six subfamilies (Tf_I, Tf_II, Tf_III, Gf_I, A_I, and A_II). As the length of the consensus 5’UTR varies among these subfamilies (34), we retained only the first two monomers plus the tether in this experiment (Fig.2D) (promoter sequences in Additional file 1: Table S1). This decision was based on two observations. First, for the L1spa element, it has been reported that a minimum of two monomers is required for detectable promoter activity (39). Second, as described earlier (Fig.1B), most of the elements from the young L1 subfamilies retain at least two intact monomers. We removed A_III subfamily from this experiment as only a small fraction of A_III elements have two intact monomers, featuring the lowest average number of monomers (Fig.1C). We incorporated two control plasmids in our dual-luciferase assays. pLK037 is a no-promoter negative control. It lacks a promoter sequence upstream of the Fluc coding sequence but contains an intact Rluc cassette; hence, its Fluc/Rluc ratio represents the assay background. To facilitate comparison of activities among different L1 promoters, we normalized the Fluc/Rluc ratio of each promoter construct to pLK037 (i.e., setting the Fluc/Rluc ratio of pLK037 to 1). pCH117 is a positive control. In Figure 2D, the normalized promoter activity for pCH117 (“L1RP”) is 914, which can be interpreted as that human L1RP 5’UTR possesses a promoter activity 914-fold above the assay background. As pCH117 usually shows the highest promoter activity among all the constructs tested, its normalized promoter activity is also an indication of the assay dynamic range. Note the assay dynamic range fluctuates to some extent from experiment to experiment (e.g., 700-to 1200-fold above background), likely due to unpredictable variations in cell status and transfection procedures. However, such fluctuations should not substantially alter the relative fold difference among promoters.

**Figure 2.**
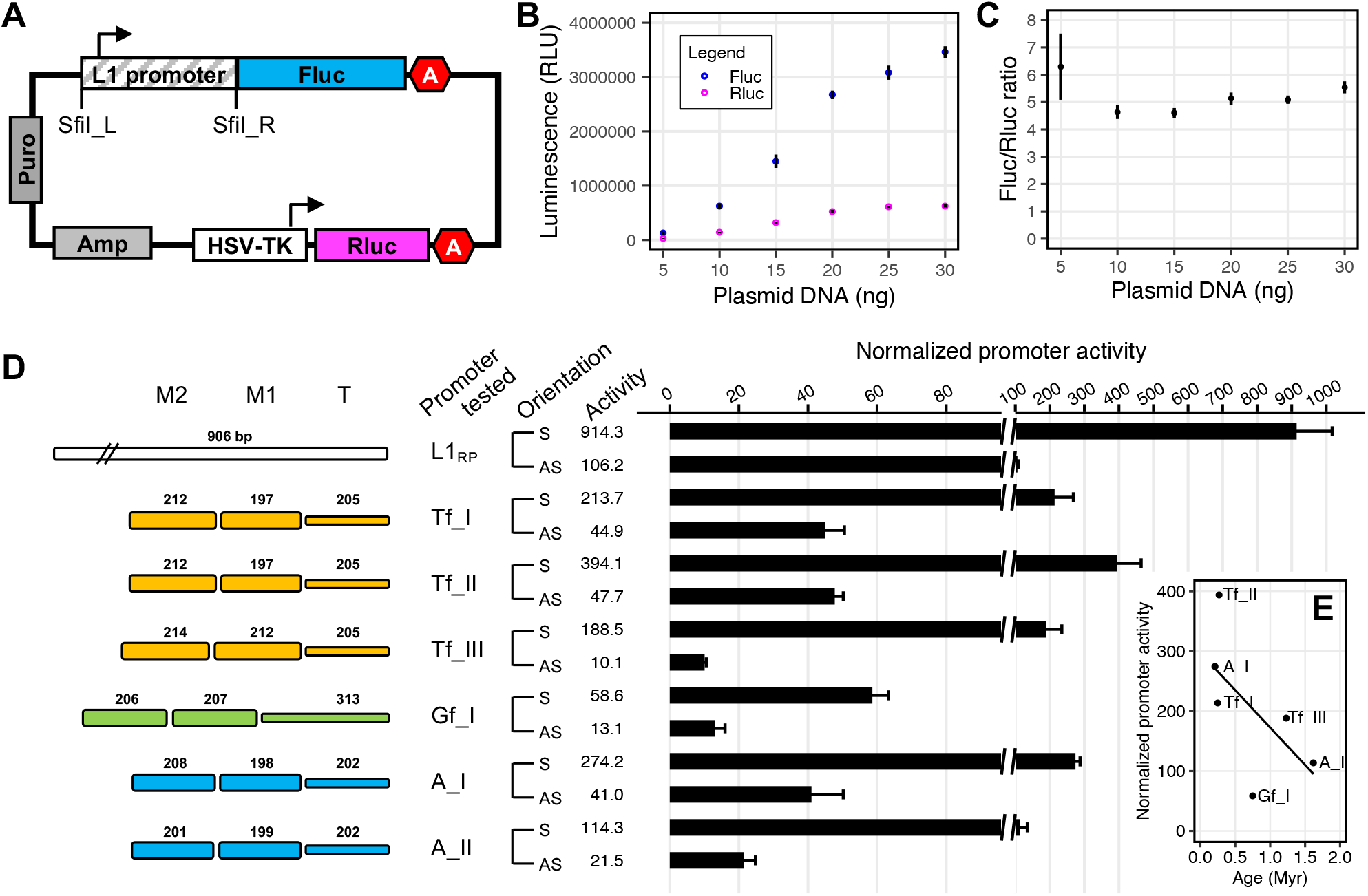
Comparison of sense and antisense promoter activities for two-monomer mouse L1 5’UTR consensus sequences. (**A**) Schematic of the dual-luciferase L1 promoter reporter assay vectors. An L1 promoter, cloned in via flanking SfiI sites, drives the firefly luciferase (Fluc) expression. A built-in Renilla luciferase (Rluc) expression cassette is used to normalize transfection efficiency. Each reporter cassette ends in a polyadenylation signal (illustrated as letter A in a hexagon). Amp, ampicillin resistance gene; HSV-TK, herpes simplex virus thymidine kinase promoter; Puro, puromycin resistance gene. Not drawn to scale. (**B**) Titration of plasmid DNA for the cell-based reporter assay. Amount of plasmid DNA is titrated in NIH/3T3 cells in quadruplicate using a control vector in which the promoter of human L1RP drives Fluc expression. The mean and standard error are shown for both Fluc and Rluc signals in raw relative luminescence units (RLU). (**C**) The calculated ratio of Fluc/Rluc from above titration experiment. Mean and standard error are shown. (**D**) Normalized activity of two-monomer consensus promoter sequences from six mouse L1 subfamilies. Sequence organization of the promoters is illustrated on the left side. The length of M2, M1, and tether (T) for each promoter is annotated (in base pairs). For each subfamily, the promoter activity was tested in both sense (S) and antisense (AS) orientation. The x-axis indicates the normalized promoter activity (i.e., the Fluc/Rluc ratio of a control no-promoter vector, pLK037, was set to 1). Note a broken x-axis is used to contrast sense and antisense promoter activities. (**E**) Inverse relationship between the sense promoter activity and subfamily age. A simple linear regression line was shown along with individual data points.

For two-monomer consensus sequences, we found the highest activity in the Tf_II subfamily (394-fold above assay background), followed by A_I (274-fold), Tf_I (214-fold), Tf_III (189-fold), A_II (114-fold), and the lowest activity in the Gf_I subfamily (59-fold). Overall, there appears to be a weak inverse relationship between subfamily age and two-monomer consensus promoter activity among these six subfamilies (simple linear regression: R = −0.62, p = 0.19) (Fig.2E). In this regard, subfamily Gf_I may be considered as an outlier, which is relatively middle-aged (0.75 MYA) but showed significantly less activity (15% of that of Tf_II).

### Differential and subfamily-dependent contribution of monomer 2, monomer 1, and tether to mouse L1 promoter activity

DeBerardinis and colleagues have previously investigated the interactions among monomers and the tether sequence based on a single promoter variant, L1spa, a prototypic mouse Tf element (38, 39). Specifically, they observed that tether alone lacked promoter activity, monomer 1 (M1) alone had some activity, either M1-T or M2 alone had about 2-fold activity above assay background, M2-M1 had about 3-fold activity, but three or more monomers showed even higher activity. These observations led to the conclusion that two monomers are required for L1 promoter activity (39). When aligned to Tf_I and Tf_II consensus sequences, L1spa showed similar levels of divergence to Tf_I and Tf_II in the 5’UTR and ORFs, but much higher similarity to Tf_I than Tf_II in the 3’UTR (e.g., all 6 SNPs are against Tf_II). Thus, we consider L1spa as a member of the Tf_I subfamily.

To validate and expand previous findings, we conducted similar studies using consensus promoter sequences for three different subfamilies, including Tf_I, A_I, and Gf_I (promoter sequences in Additional file 1: Table S2). For Tf_I subfamily (Fig.3A), consistent with the previous report using L1spa 5’UTR (39), the promoter construct with two tandem monomers and the tether (M2-M1-T) showed 6.0-fold higher activity than the construct containing M1 and the tether (M1-T). The previous study showed minimal activity from tether alone or M1 alone, but M2 alone was not tested. The wide dynamic range of our assay allowed us to differentiate the relative activities of M2, M1, and tether. In the context of the consensus sequence, M2 alone displayed an activity equivalent to 22.2% of the M2-M1-T sequence. M1 alone is about 2-fold less active (13.0% of M2-M1-T) but remains 12-fold above the assay background. Tether alone showed even less activity (4.1% of M2-M1-T) but remained 11.6-fold above the assay background (Welch t-test, p = 0.002). To confirm such residual promoter activities, we included two additional control plasmids (Fig.3A). First, we replaced the promoter sequence with a 205-bp fragment from the green fluorescent protein (GFP) coding sequence, equivalent to the length of Tf_I tether. As expected, this 205-bp GFP (GFP205) sequence showed no promoter activity (0.6-fold relative to the assay background). Second, we placed the tether sequence in its antisense orientation (T_AS). Interestingly, the antisense Tf_I tether had 8.2-fold higher activity than the assay background (Welch t-test, p < 0.001). These results suggest that the Tf_I tether sequence has some weak transcriptional activities in both sense and antisense orientations. To aid in the interpretation of the contribution of individual domains, we diagrammed promoter activities along with domain locations in an integrated manner (Fig.3B). For Tf_I subfamily, M2-M1-T has the highest activity, 3.2-fold higher than any other permutations of its subdomains. Comparing M1-T with T and M1, it seems that the activity of M1-T is the sum of M1 and T alone, suggesting an additive role. The addition of M2 to M1-T appears to be synergistic, as the resulting M2-M1-T construct is 6-fold higher than M1-T. To probe the contribution of M1 to overall two-monomer promoter activity, we generated a synthetic construct in which M2 is directly placed upstream of the tether (M2-T) (Fig.3A). Comparing M2-T with M2-M1-T, the deletion of M1 reduced the promoter activity by at least 3-fold. This result suggests that M1 positively contributes to the 2-monomer promoter activity for Tf_I subfamily. Taken together, all three domains contribute positively to the overall two-monomer 5’UTR activity in Tf_I subfamily.

For A_I subfamily, M2-M1-T displayed 30.4-fold higher activity than M1-T (Fig.3C). The reduction is even more dramatic than that observed for the Tf_I subfamily. Then we examined the activities of each domain: M2, M1, and tether alone. Surprisingly, the A_I M2 showed remarkable promoter activity on its own, with 3.6-fold higher activity than the two-monomer construct. In contrast, M1 and tether had low but detectable amount of activity relative to the assay background. Specifically, both had less than 3% of M2-M1-T but still 7∼8-fold above the assay background (Welch t-test, p = 0.002 for M1 and p < 0.001 for T). However, combining M1 and T together did not lead to any substantial increase in promoter activity (10-fold above background for M1-T). The deletion of M1 from M2-M1-T reduced the promoter activity by a mere 7% (comparing M2-T with M2-M1-T), suggesting M1 contributes little to the overall two-monomer promoter. On the other hand, the presence of tether sequence reduced M2 activity by 4-fold (comparing M2 and M2-T), indicating that A_I tether significantly suppresses the promoter activity of M2 and likely plays a negative role in the context of two-monomer promoter. Thus, M2 dominates in its contribution to the overall A_I promoter activity. Similar to the experiment with Tf_I promoters, a 202-bp fragment from the GFP coding sequence (GFP202), equivalent to the length of A_I tether, showed little promoter activity (1.5-fold above background). The antisense A_I tether had 3-fold higher activity than the assay background (Welch t-test, p < 0.001). These results suggest that the A_I tether sequence also has some weak transcriptional activities in both sense and antisense orientations. To summarize, M2 is the major contributor of two-monomer promoter activity for A_I subfamily, the tether negatively regulates M2 activity in the context of two-monomer 5’UTR, while the role of M1 is minimal (Fig.3D).

Similar trend was observed for Gf_I promoter (Fig.3C). Gf elements were first described in 2001 by Goodier and colleagues (35). The Gf_I subfamily (34) conforms to pattern II of Gf promoters in the original scheme. As described earlier, the consensus Gf_I M2-M1-T construct had much weaker promoter activity than the corresponding Tf_I and A_I constructs (27.4% and 21.4%, respectively; Fig.2D). Nevertheless, it remained 3.2-fold more active than M1-T (Welch t-test, p < 0.001), although the magnitude of reduction was not as dramatic as in A_I and Tf_I. The activities of individual domains, M2, M1 and the 313-bp tether, were 20.2%, 9.8%, and 20.4% of M2-M1-T, respectively, but remain significantly above the assay background (Welch t-test, p = 0.003, 0.002 and 0.0002, respectively). The antisense 313-bp tether (T_AS) also had substantial amount of promoter activity (26.6% of M2-M1-T; Welch t-test, p = 0.001 against the assay background). Note the 313-bp tether includes a truncated 64-bp monomer at its 5’ end. We also subcloned the tether sequence without the 64-bp truncated monomer. The shortened 249-bp tether had detectable activities in both sense (T249, 11.8% of the 2-monomer promoter; Welch t-test, p = 0.03 against assay background) and antisense orientation (T249_AS, 13.7% of 2-monomer promoter; Welch t-test, p < 0.001 against assay background). The interactions among individual domains for subfamily Gf_I are distinctly different from both Tf_I and A_I (Fig. 3F). For Gf_I, the interaction between M1 and T appears to be additive when comparing M1-T with M1 and T alone. On the other hand, M2 and M1-T are somewhat synergistic as M2-M1-T is about 2-fold the sum of M2 and M1-T. In comparison, the deletion of M1 only reduced the promoter activity for Gf_I by 13%, suggesting M1 plays a minor role in Gf_I subfamily. Thus, the two-monomer activity of Gf_I is mainly the result of interaction between M2 and tether.

**Figure 3.**
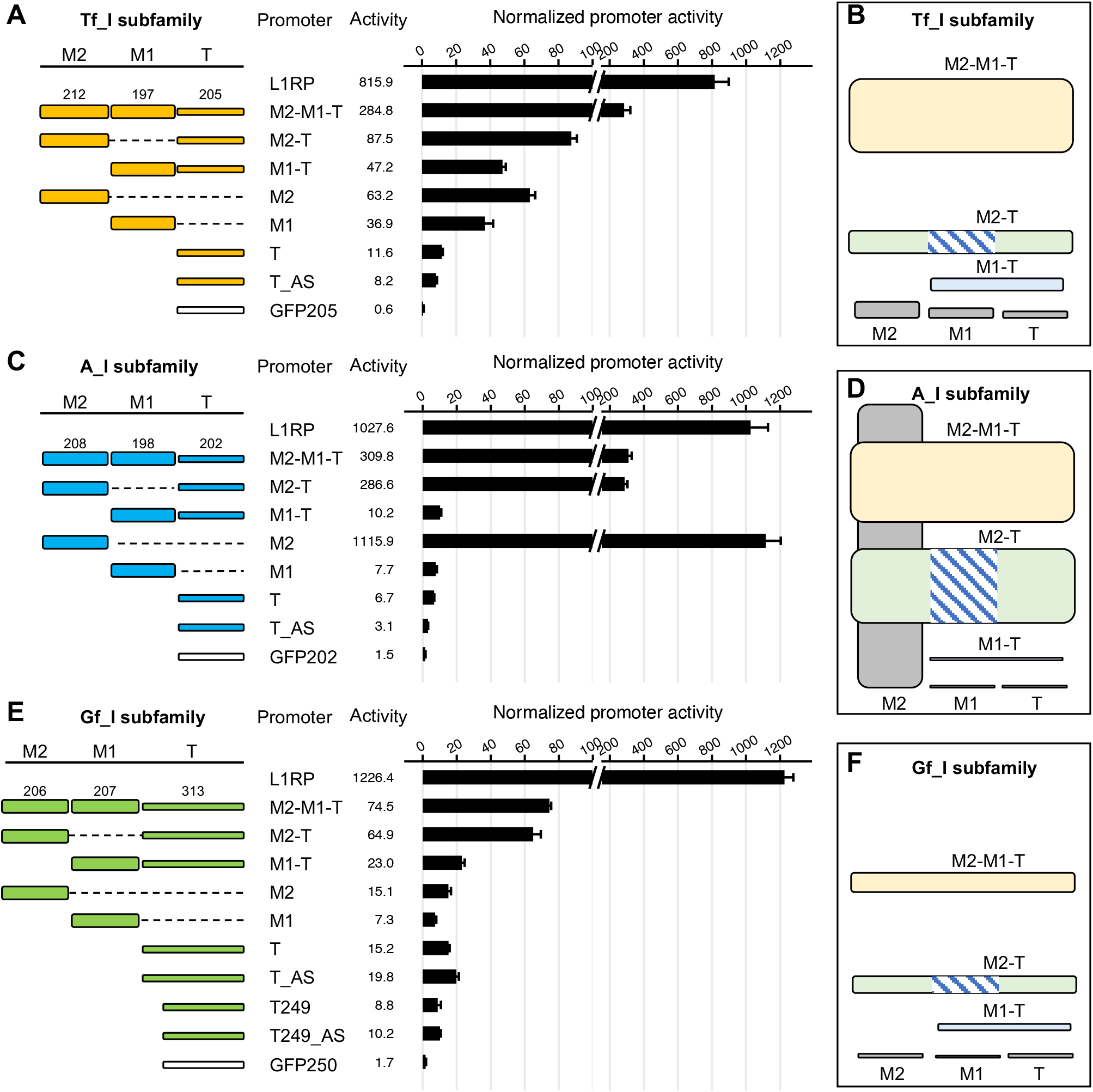
Differential contribution of monomer 2, monomer 1 and tether to overall promoter activity. Normalized promoter activity of individual 5’UTR domains for subfamily Tf_I (**A**), A_I (**C**), and Gf_I (**E**). Sequence organization of the promoters is illustrated on the left side. The length of M2, M1, and tether for each promoter is annotated (in base pairs). The dashed line represents domain(s) that were removed in reference to the two-monomer 5’UTR sequence (M2-M1-T). The tether was tested in both sense (T) and antisense (T_AS) orientation. A short version of Gf_I tether was additionally included (T249 and T249_AS) in panel E. The x-axis indicates the normalized promoter activity (i.e., the Fluc/Rluc ratio of a control no-promoter vector, pLK037, was set to 1). Note a broken x-axis was used to highlight the wide range of promoter activities. On the right hand are 2-D representations of the promoter data for subfamily Tf_I (**B**), A_I (**D**), and Gf_I (**F**), corresponding to panel A, panel C, and panel E, respectively. Each domain tested is represented by a filled box. The domains are arranged in the order of M2, M1, and tether from left to right. The height of the box corresponds to the normalized promoter activity (to scale). The hatched lines represent the missing M1 domain in the M2-T promoter construct.

### Length of monomer 3 has a U-shaped effect on overall promoter activity

Thus far, we have shown the contribution of individual M2, M1, and T sequences in the context of a two-monomer 5’UTR for Tf_I, A_I, and Gf_I subfamilies. However, many L1 promoters contain more than two monomers. Indeed, for the two youngest mouse L1 subfamilies, Tf_I and A_I, more L1 promoters start in M3 than in any other positions (157 out of 513 or 30.6%, and 357 out of 1125 or 31.7%, respectively) (Fig. 1B). On the other hand, the distribution of the 5’ start positions in M3 is, albeit varied, nonrandom. For example, 16.6% (26/157) of the Tf_I loci containing M3 start at nucleotide position 83 (Fig. 4A) and 26.3% (94/357) of the A_I loci containing M3 start at nucleotide position 86 (Fig. 4B). To dissect the role of varied lengths of monomer 3, we conducted a direct comparison between M3-M2-M1-T and M2-M1-T for both Tf_I and A_I subfamilies (Fig.4C-D) (promoter sequences in Additional file 1: Table S3). Indeed, both three-monomer consensus constructs were more active than the two-monomer counterparts. For Tf_I subfamily, the three-monomer promoter was 2.4-fold higher than the two-monomer version and was only 17.4% lower than the reference L1RP promoter (Fig.4C). For A_I subfamily, the three-monomer promoter was 4.0-fold higher than the two-monomer version and even outperformed the highly active L1RP promoter by 19.3% (Fig.4D). To study the impact of an incomplete monomer on the overall promoter activity, we created series of A_I and Tf_I promoter constructs by truncating the third monomer stepwise for 40 bp. For Tf_I subfamily, the deletion of the first 40 bp reduced the promoter activity to 74.0% of the three-monomer construct (Fig.4C). The removal of the first 80 bp reduced the promoter activity further to 36.5% of the three-monomer construct. Deletion of the first 122 bp had additional effect (down to 23.6% of the three-monomer construct). However, this diminishing trend was reversed when the promoter was further truncated. The promoter activity was restored to 31.6% of the three-monomer construct when the first 162 bp was deleted. The deletion of the entire third monomer (212 bp), giving rise to the two-monomer construct, restored the activity to 42.3% of the three-monomer construct. Similar patterns were seen with the vector series for A_I subfamily (Fig.4D). The promoter activity was reduced to 45.6%, 18.0%, 15.7% of the three-monomer construct with 40-, 80-, 122-bp deletions, respectively, and then rebounded back to 18.1% and 25.3% of the promoter activity with deletion of 160 bp and the entire 208-bp M3, respectively. Thus, for both subfamilies, the first 80 bp of M3 has a positive impact on overall promoter activity but the last 80 bp negatively regulates the promoter activity. The interaction between the length of M3 and the overall promoter activity is characteristic of an asymmetrical U-shaped relationship (Fig.4C-D).

**Figure 4.**
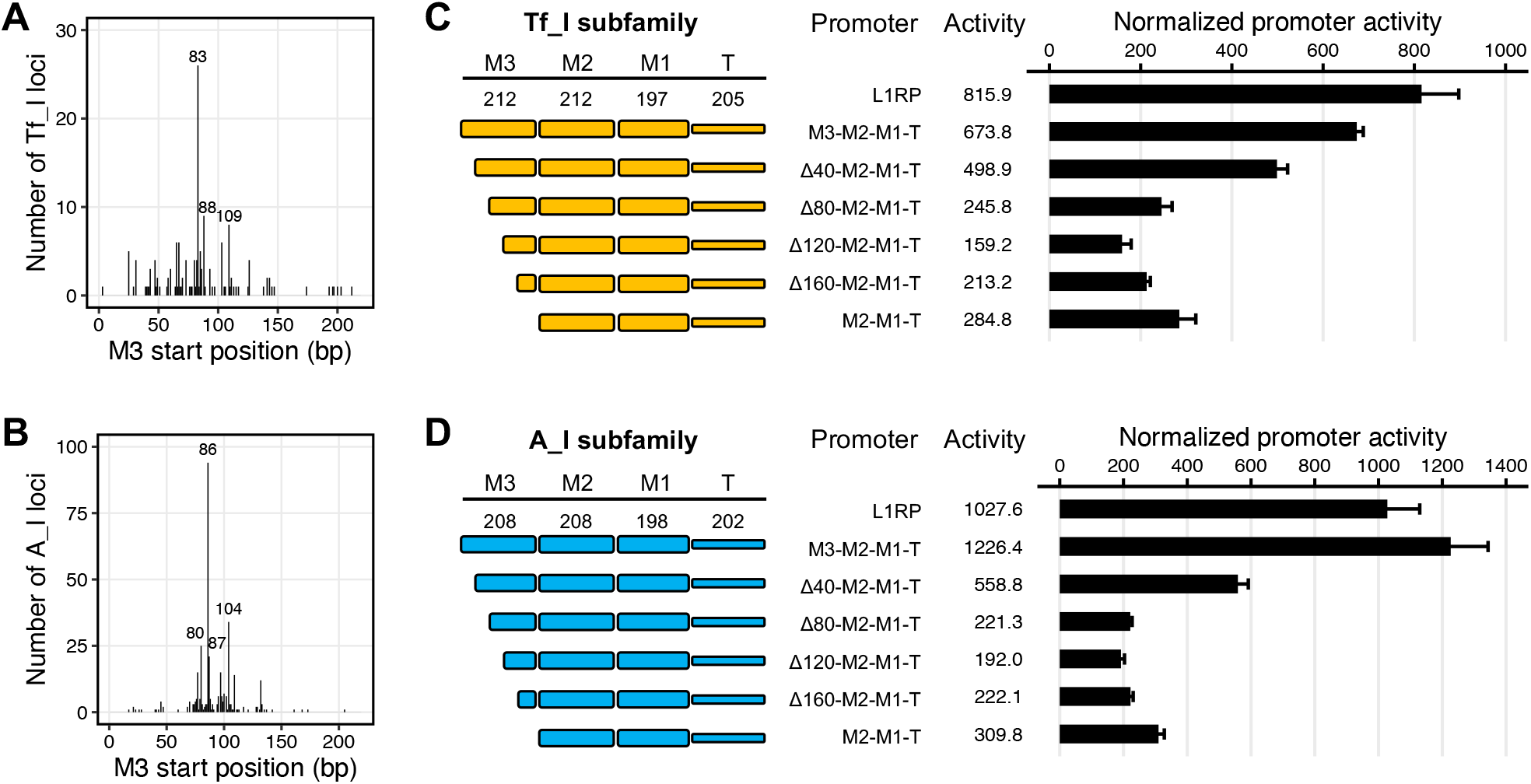
Contribution of different lengths of monomer 3 to overall promoter activity. (**A**) Distribution of 5’UTR start positions for 157 Tf_I loci that are 5’ truncated within M3. The x-axis represents nucleotide positions from 1 to 212 for M3. The y-axis displays the count of Tf_I loci that start at each nucleotide position. The three most frequent nucleotide positions are annotated. (**B**) Distribution of 5’UTR start positions for 357 A_I loci that are 5’ truncated within M3. (**C**) Normalized promoter activity of Tf_I 5’UTR consensus sequences with varying M3 length. Sequence organization of the promoters is illustrated on the left side. The length of M2, M1, and T for each promoter is annotated (in base pairs). The x-axis indicates the normalized promoter activity. Note promoter constructs in panel C were tested together in the same 96-well plate with those for Fig.3A; thus, L1RP, M3-M2-M1-T and M2-M1-T are shared between Fig.3A and Fig.4C. (**D**) Normalized promoter activity of A_I 5’UTR consensus sequences with varying M3 length. Note promoter constructs in panel D were tested together with those for Fig.3B.

### Two-monomer consensus sequences have antisense promoter activities

The human L1 contains an antisense promoter activity (27), which affects as many as 4% of the human genes (43). An antisense promoter activity has been previously reported in ORF1 region of the mouse L1 (44). However, it remains unclear whether mouse L1 5’UTRs have antisense promoter activities. To uncover potential antisense promoter activities, we inverted the two-monomer consensus sequences from the six young mouse L1 subfamilies and compared them to their sense-oriented counterparts (Fig.2D). In our control experiment, the antisense oriented L1RP 5’UTR showed 106.2-fold activity above the experimental background, equivalent to 11.6% of that of the sense promoter. The relative strength of antisense versus sense promoter activity for L1RP reported here is consistent with a previous report (45), which showed 8-fold lower activity for the antisense promoter than the sense promoter by both Northern blot and a luciferase-based reporter assay in human HeLa cells. In this context, all six L1 subfamilies demonstrated detectable levels of antisense promoter activities (Fig.2D). The three youngest subfamilies (A_I, Tf_I, and Tf_II) all had >40-fold activity above the assay background in the antisense orientation, equivalent to 15.0%, 21.0%, and 12.1% of the activity from the corresponding sense promoter, respectively. The antisense sequence of A_II subfamily showed 21.5-fold activity in the reporter assay, which is equivalent to 18.8% of the sense promoter. Gf_I and Tf_III subfamilies had the lowest antisense promoter activities (13.1 and 10.1-fold above assay background, respectively), corresponding to 22.3% and 5.3% of their sense promoter counterparts.

## Discussion

The two-monomer 5’UTRs tested in this study are consensus sequences as defined by the Boissinot group in 2013 (34). For subfamilies with recent periods of activity, it is expected that individual copies be similar to the consensus sequence (46). Indeed, this prediction is true for the three youngest subfamilies (A_I, Tf_I, and Tf_II; Additional file 1: Table S4). The reference mouse genome contains 21 identical loci and 134 single-mismatch loci for the 608-bp A_I two-monomer 5’UTR sequence, three identical loci and 33 single-mismatch loci for the 614-bp Tf_I two-monomer sequence, and 18 single-mismatch loci for Tf_II two-monomer sequence. In contrast, for the middle-aged Gf_I subfamily, only three single-mismatch loci are found for its 726-bp two-monomer 5’UTR sequence. The older Tf_III and A_II subfamilies do not have any loci carrying less than three mismatches. Therefore, our results not only reflect the promoter activities of the consensus 5’UTR sequences tested but can potentially be extended to a number of endogenous mouse L1 loci, especially for A_I, Tf_I, Tf_II, and Gf_I.

In the context of two-monomer 5’UTRs, the inclusion of M2 upstream of M1 is essential for its enhanced promoter activity. The enhancement by M2 is 6.0-fold for Tf_I, 30.4-fold for A_I, and 3.2-fold for Gf_I (Fig.3; comparing M2-M1-T with M1-T for each subfamily). When normalized to the control L1RP promoter, it is evident that the activity of A_I M2 consensus (108.6% of L1RP) far exceeds that of Tf_I (7.7% of L1RP) and Gf_I (1.2% of L1RP) in NIH/3T3 cells (Fig.3). Note the definition of individual monomers is not necessarily consistent in the literature across mouse L1 subfamilies. As expected, sequence alignment shows extensive sequence divergence among A_I, Tf_I, and Gf_I M2 sequences used in this study (Additional file 2: Fig.S1). For the 208-bp A_I M2 consensus sequence (5’-GTGCCTGCCC&#8230;GTGGAACACA-3’), we defined its boundary in the A_I 5’UTR consensus sequence by following the convention established by Loeb and colleagues when type A monomer was first described (11) (Additional file 2: Fig.S2). Comparing with previously described A monomer consensus sequences (41, 47), the A_I M2 sequence has three mismatches. BLAST search of this A_I M2 sequence in the mm10 mouse genome assembly returns 67 identical hits and 138 single-mismatch hits (Additional file 1: Table S4). Coincidentally, this A_I M2 sequence is identical to the A monomer subtype 1 recently defined by the Smitch group using a profile-HMM based unsupervised approach (48). For the 212-bp Tf_I M2 consensus sequence (5’-GACAGCCGGC…GTGGGCCGGG-3’), we followed the convention initially established the Kazazian group (38, 39) (Additional file 2: Fig.S3). It differs from Naas’s version (38) by one nucleotide at position 171 and from DeBerardinis’s version (39) by an additional nucleotide at position 24. Seventeen copies identical to the consensus Tf_I M2 sequence are present in the mouse genome (Additional file 1: Table S4). Note the T monomers recently identified by the profile-HMM approach would start at nt 135 (5’-GGTGCGCCAG…-3’) (48). The 212-bp Tf_I M2 tested here displays a single mismatch with T monomer subtype 22 at nt 24 and with subtype 25 at nt 102, respectively. The 206-bp Gf_I M2 consensus sequence (5’-TGAGAGCACG…ACCTTCCTGG-3’) follows the original boundary definition but differs from Goodier’s version by two nucleotides at nts 152-153 (35) (Additional file 2: Fig.S4). It has 121 identical copies in the mouse genome (Additional file 1: Table S4). Note the Gf monomer subtype 2 defined by the profile-HMM approach (48) would start at position 204 but is otherwise identical to the Gf_I M2 sequence tested in this study. How individual SNPs affect each monomer variant’s activity necessitates future studies.

Our study highlights the difference between M2 and M1 in promoter activity. The most dramatic example is from the A_I subfamily. In head-to-head comparison, its M1 alone has a mere 7.7-fold activity above assay background but its M2 is 145-fold more active than M1 (Fig.3B). This functional difference reflects the sequence divergence between them. The A_I M2 and M1 are 86.5% (180 out of 208 nucleotides) identical (Additional file 2: Fig.S2). Besides 18 SNPs, M1 possesses three short deletions, including the deletion of one copy of the tandem ACTCGAG motif noted previously (48). For Tf_I subfamily, the M2 and M1 are 76.6% (164/214) identical overall (Additional file 2: Fig.S3). The divergence is concentrated in the second half of the monomers, with the putative YY1 binding motif preserved in M1. Despite the larger difference than seen in subfamily A_I, Tf_I’s M2 and M1 only differed in promoter activity by 1.7-fold (Fig.3A). For subfamily Gf_I, its M2 and M1 are highly similar with 96.6% identity (200/207) (Additional file 2: Fig.S4). The seven mismatches are located toward the 3’ end of the sequence. At the functional level, M2 is 2-fold more active than M1 (Fig.3C). Future studies are necessary to pinpoint the key nucleotide positions that are responsible for differential promoter activity between these M2 and M1 sequences. It should also be noted that, while our study focused on a few consensus monomers, the mouse genome contains a large number of A or Tf monomer subtypes, which display different modes of position preference within a 5’UTR monomer array (48). It is entirely possible that a strong monomer, similar to A_I M2, is positioned directly upstream of a tether, forming a highly active one-monomer-tether 5’UTR. Therefore, one could not automatically assume low promoter activity for a shortened M1-T like locus.

Unlike monomer sequences, the tether sequences share a significant amount of homology among the three subfamilies (Additional file 2: Fig.S5). The tethers for subfamily A_I and Tf_I are similar in length and 76.6% identical. Both have modest activities (7.7-fold or 11.6-fold above assay background, respectively) (Fig.3A-B). For subfamily Gf_I, two different versions of tether were tested. One is 249 bp long, which can be divided into a 3’ 208-bp segment (with 84.1% identity to Tf_I tether) and a 5’ 41-bp segment (equivalent to 5’ extension into the corresponding Tf_I M1 region). It showed 8.8-fold activity above assay background (Fig.3C). The other is 313 bp long. The addition of the extra 64 bp truncated Gf_I monomer rendered the longer tether sequence slightly more active (15.2-fold above assay background). Despite the modest activity on its own, the tether sequence seems always augment the activity from M2 or M1 to some extent. The only exception is when it is coupled with A_I M2 as described earlier. The molecular mechanism via which the tether contributes to the overall promoter activity is unknown. The high level of sequence conservation among all A, Tf, Gf and F subfamilies reflects its common ancestry (34). Though highly speculative it is possible that the tether region has other regulatory roles during L1 replication cycle.

We demonstrated antisense promoter activity for two-monomer 5’UTR constructs from all seven evolutionarily young mouse L1 subfamilies examined (Fig.2D). The amount of antisense promoter activity is a fraction of the corresponding sense promoter activity, ranging from 5% to 22%. Notably, when tested in multiple cell lines, the antisense promoter activity of human L1PA1 5’UTR falls within this range (11.6% in NIH/3T3 cell line [this study], 12.5% in HeLa cell line (45), 7.8% in human embryonal carcinoma 2102Ep cell line (29), and 25% to 33% in human embryonic stem cell lines (29)). The relative contribution of M2, M1, and tether domains to the overall antisense promoter activity remains unclear. When the tether sequence from subfamily Tf_I, A_I, and Gf_I was tested in the antisense orientation, it showed 2.9%, 1%, and 26.5% of the corresponding two-monomer promoter, respectively (Fig.3), suggesting only Gf_I tether contributes substantially to the antisense promoter activity. Our findings on antisense promoter activity in mouse L1 5’UTRs contract with a previous study, which found minimal activity for two individual A type monomers and a tether sequence when tested in the antisense orientation (41). This discrepancy may be explained by differences in the sensitivity and dynamic range of the reporter assays used and the promoter sequences tested. On the other hand, our results are consistent with a recent analysis of cap analysis of gene expression (CAGE) data from mouse embryonic testes, showing strong antisense transcription start site (TSS) signals for Gf and T monomers (48).

In reference to the computationally defined monomers, the 5’ termini of endogenous L1 loci display a tendency of starting from certain nucleotide positions. The 5’ truncation points of Tf monomers, including the two prototypic full-length Tf insertions, are clustered at nts 70-110 (38, 39, 48). This region overlaps with a putative YY1 binding motif GCCATCTT at nts 80-87, which has been postulated to play a similar function in controlling transcription initiation as reported for human L1 5’UTR (39, 48, 49). Earlier observations from a limited number of A type loci indicated two clusters of 5’ truncation points relative to a complete monomer (two monomers start at nts 24-25 and ten start at nts 70-85) (11, 47, 50). A recent genome-wide analysis confirmed the predominance of truncation points within a 30-bp region at nts 70-100 for the 5’ most A monomers (48). Notably, a tandem ACTCGAG motif of unknown function is present at nt 98-111 (36, 48). Our own analysis at single-base resolution replicated these findings, showing a broader distribution with a dominant peak at nt 83 for Tf_I monomers (Fig.4A) and a much tighter distribution with a dominant peak at nt 85 for A_I monomers (Fig.4B). However, the role of a partial or incomplete monomer at the beginning of a mouse L1 5’UTR had not been addressed by previous studies. Using the consensus A_I and Tf_I 5’UTR as a model, we found a complex U-shaped relationship between the length of the outer M3 and the overall promoter activity (Fig.4C-D). As expected, promoters with three full monomers are much more active than those with two monomers for both subfamilies. However, the lowest promoter activities were found when 122 bp (but not when additional sequences) was removed from the 5’ end of the M3. Thus, the contribution of M3 sequence to overall promoter activity is not simply proportional to its length. This phenomenon is consistent with a model in which both M3 and its downstream monomers promote parallel transcription initiation events (11). Under this model, the deletion of 122 bp from M3 abolishes transcription initiation from M3 and unmasks negative regulation of transcription initiation from M2 by the remaining M3 sequence, leading to much reduced overall transcription output. Addition deletion of M3 sequence eliminates the negative regulation and enables unimpeded transcription initiation from M2. The consensus M3 and M2 sequences are not identical though: they differ by two nucleotides in A_I (Additional file 2: Fig.S2), and by three nucleotides in Tf_I (Additional file 2: Fig.3). Nevertheless, according to the distribution of the 5’ start positions of endogenous loci that are 5’ truncated within M3 (Fig. 4A-B), one would predict that most of such Tf_I and A_I elements be transcribed at lower levels than an element with either three or two full-length monomers. This observation raises an interesting question about the molecular processes leading to such a 5’ truncation pattern and any advantages or disadvantages toward subsequent rounds of L1 replication.

## Conclusions

The multimeric nature of mouse L1 5’UTRs presents a challenge to investigate mouse L1 transcriptional regulation. Accordingly, unlike the human L1 5’UTR, many aspects of mouse L1 transcription remain poorly understood. In this study, aided by synthetic biology and report assays with a wide dynamic range, we compared sense promoter activities and discovered antisense promoter activities from six evolutionarily young mouse L1 subfamilies. Expanding upon a pioneering study featuring a single Tf_I element, we determined contribution of monomer and tether sequences among three main lineages of evolutionarily young mouse L1s: A_I, Tf_I and Gf_I. Our work validated that, across multiple subfamilies, having the second monomer is always much more active than the corresponding one-monomer construct. For individual promoter components (M2, M1, and tether), M2 is consistently more active than the corresponding M1 and/or the tether for each subfamily. More importantly, we revealed intricate interactions between M2, M1 and tether domains and such interactions are subfamily specific. Using three-monomer 5’UTRs as a model, we established a complex nonlinear relationship between the length of the outmost monomer and the overall promoter activity. Overall, our work represents an important step toward elucidating the molecular mechanism of mouse L1 transcriptional regulation and L1’s impact on development and disease.

## Materials and Methods

### Computational analysis of mouse L1 5’UTR start positions

BLAST+, a suite of command-line tools to run BLAST locally (51), was used to search for the promoter region (query sequence) in each L1 sequence (subject sequence). For each subfamily, we created a query sequence containing 11 monomers and the corresponding tether sequence by removing the 5’ partial monomer from the consensus sequence (34) and appending copies of the last full-length monomer to the 5’ end of the consensus sequence until there was a total of 11 monomers. The monomers duplicated in the 11-monomer query sequences were the 212-bp M3 for Tf_I and Tf_II, the 214-bp M3 for Tf_III, and the 208-bp M3 for A_I, A_II and A_III. We derived four separate 11-monomer query sequences for Gf_I, corresponding to the four 5’UTR monomer organization patterns defined previously (35). However, pattern III was later excluded from downstream analyses since nearly all its alignments were short and overlapped with alignments with other patterns. Patterns I, II and IV differ from each other in tether length (377, 313, and 250 bp, respectively). Pattern II is considered as a prototype for Gf_I; its 206-bp M2 was duplicated to make the 11-monomer query. The same M2 was used to populate all monomer positions for patterns I and IV. L1 sequences belonging to subfamilies Tf_I, Tf_II, Tf_III, Gf_I, A_I, A_II and A_III were extracted from the mouse genome assembly GRCm38/mm10 using SeqTailor (52), and saved as subfamily-specific subject sequence files. The input BED files containing genomic coordinates for individual L1 loci were derived from mm10 Repeat Library db20140131, which is available from the RepeatMasker website (53). For each subfamily, the query sequence was searched against each subject sequence in the subject sequence file using BLAST+. The parameters used were “-perc_identity 0, -num_threads 4, -max_target_seqs n” (where n is a number greater than the total number of sequences in the local database). The output alignment file was then parsed in RStudio with R version 3.6. We filtered out alignments that do not end in the last 10 bases of the corresponding tether region of the query sequence and alignments that do not start within the first 10 bases of the subject L1 sequence. This filtering step removed potential loci with a 3’ truncated tether and/or with a chimeric 5’UTR composed of monomers from divergent L1 subfamilies. For Gf_I, five loci were shared between patterns I and II, and three of them were also shared with pattern IV. The redundant entries were removed, and the five loci were retained under pattern II only. To plot the 5’ start position of L1 sequences in reference to the monomer or tether positions in the query sequence, the start of the alignment in query was separated into 12 bins (tether, and M1 to M11; see Fig.1B). To calculate the average number of monomers for each subfamily, we excluded the small number of loci that start either in the tether or M11+ (see Fig.1C). The 5’ start position of each locus relative to the specific monomer position in the query was used to determine the factional length of the 5’UTR. The copy number of two-monomer promoters and individual monomer and tether domains in the mouse genome (see Additional file 1: Table S4) was determined in a similar fashion using BLAST+.

### Plasmid construction

A detailed list of the promoter constructs, including primers and the corresponding promoter sequences, is provided as supplemental tables (Additional file 1). pCH036 is the base vector for inserting individual promoter sequences between two heterotypic SfiI sites (Fig.2A; SfiI_L=GGCCAAAA/TGGCC and SfiI_R=GGCCTGTC/AGGCC; “/” indicates the cleavage site) immediately upstream of the Fluc reporter gene. It looks nearly identical to all the derivative dual luciferase assay vectors except the “L1 promoter” sequence is substituted by a 48-bp multiple cloning site segment. Originating from pESD202, the double-SfiI cassette enables directional inert swapping via a single, robust restriction/ligation cycle (54). We derived pCH036 from pLK003. The latter was similar in vector architecture to pCH036 but, instead of the Fluc reporter gene, pLK003 had a firefly luciferase based retrotransposition indicator cassette (FlucAI). To make pCH036, we amplified the Fluc reporter gene from pGL4.13 (Promega) using PCR primers WA1312 5’-AAAACCTAGGGGCCTGTCAGGCCATGGAAGATGCCAAAAACATTAAGAAG-3’ and WA1314

5’-AAAAGGTACCTTACACGGCGATCTTGCCG-3’. The backbone fragment of pLK003 was prepared by a double digestion with AvrII and KpnI, removing the FlucAI cassette, and subsequently ligated to the Fluc PCR fragment with the same sticky ends. In the resulting pCH036, the second SfiI site (i.e., Sfil_R) is immediately upstream of the start codon of Fluc.

pCH117 is a positive control vector that contains the human L1RP 5’UTR as the “L1 promoter”. To make pCH117, we amplified the L1RP 5’UTR from pYX014 (55). The PCR product was digested with SfiI (New England Biolabs), gel purified, and ligated with SfiI-digested pCH036. pLK037 is a negative control vector that contains an empty double-SfiI cassette upstream of the Fluc reporter gene. It was derived by SfiI digestion of pCH117, blunting of the 3’ overhangs with Klenow fragment of E. coli DNA polymerase I (New England Biolabs), and self-ligation of the backbone fragment. pLK043, pLK044, and pLK045 are control vectors that contain 202-, 205-, and 250-bp of EGFP coding sequence in the double-SfiI cassette, respectively. The corresponding EGFP sequences were amplified from pWA003 (55) by using the same reverse primer paired with three different forward primers. The PCR product was digested with SfiI, gel purified, and ligated with SfiI-digested pCH036.

The three-monomer Tf_I consensus promoter in pLK086 was derived from a synthetic DNA fragment that is flanked by SfiI_L and Sfil_R restriction sites. Primers were designed to serially truncating M3 by 40-, 80-, 122-, and 162-bp from the 5’ end. The resulting PCR products were SfiI digested and ligated into SfiI-digested pCH036, giving rise to pLK094, pLK095, pLK096, and pLK097. The two-monomer Tf_I promoter in pLK050 was derived from a synthetic DNA fragment. Primers were designed to amplify M2, M1, and T. The resulting PCR products were digested and ligated into pCH036, resulting in pLK057, pLK056, and pLK054. The antisense version of the tether fragment was similarly cloned into pLK055. M2-T sequence in pLK098 and M1-T sequence in pLK047 were derived from synthetic DNA fragments.

The three-monomer A_I consensus promoter in pLK085 was derived from a synthetic DNA fragment. Primers were designed to serially truncating M3 by 40-, 80-, 122-, and 160-bp from the 5’ end. The resulting PCR products were SfiI digested and ligated into SfiI-digested pCH036, giving rise to pLK090, pLK091, pLK092, and pLK093. The two-monomer A_I promoter in pLK049 was derived from a synthetic DNA fragment. Primers were designed to amplify M2, M1, M1-T and T. The resulting PCR products were digested and ligated into pCH036, resulting in pLK053, pLK052, pLK040 and pLK041. The antisense version of the tether fragment was similarly cloned into pLK042. M2-T sequence in pLK046 was derived from a synthetic DNA fragment.

The two-monomer G_I consensus promoter in pLK051, the M2-T promoter in pLK099, the M1-T promoter in pLK048 were derived from separate synthetic DNA fragments. Primers were designed to amplify M2 and M1, respectively. The resulting PCR products were digested and ligated into pCH036, resulting in pLK063 and pLK062. Two different lengths of tether were considered. Primers were designed to amplify and clone the tether as a 313 bp fragment in either sense (pLK060) or antisense orientation (pLK061). A shortened 249 bp version of the tether was also cloned in either sense (pLK058) or antisense (pLK059) orientations.

The two-monomer consensus promoters for A_II (pLK087), Tf_II (pLK088), and Tf_III (pLK089) were derived from separate synthetic DNA fragments. All synthetic DNA fragments were purchased from either Genewiz (part of Azenta Life Sciences) or Twist Biosciences. pJT01, pJT02, pJT03, pJT04, pJT05, pJT06, and pJT07 contain antisense versions of the 2-monomer promoters in pLK049, pLK050, pLK051, pLK087, pLK088, pLK089 and of the L1RP promoter in pCH117, respectively. To make these antisense promoter constructs, primers were designed to amplify the sense-oriented promoters from the respective precursor constructs so resulting PCR fragments would reverse the orientation of the promoter with respect to the two heterotypic SfiI sites.

### Cell line authentication

All promoter assays were performed in a subline of NIH/3T3 mouse embryonic fibroblast cells maintained in our lab. To confirm cell identity, we submitted an aliquot of the cells to American Type Culture Collection (ATCC) for mouse short tandem repeat (STR) testing. The testing involved the analysis of 18 mouse STR loci as well as two specific markers to screen for potential cell line contamination by human or African green monkey species (56). The STR profile of our cells is nearly identical to the ATCC reference NIH/3T3 cell line (ATCC CRL-1658). Specifically, our subline shares all 26 alleles that are present in ATCC NIH/3T3 at the 18 mouse STR loci analyzed. In addition, it has evolved a second allele at the STR locus 6-4 (the new allele is one repeat longer than the reference allele). The complete cell line authentication report is available as a supplemental document (Additional file 2: Fig.S6).

### Dual-luciferase promoter assay

Assays were performed with NIH/3T3 cells in 96-well format. Cells were first trypsinized from a stock dish, diluted into a suspension at 200,000 cells per ml, and kept at 37°C before seeding into a 96-well plate. Lipofectamine 3000 (Invitrogen) was used following a reverse transfection protocol. Briefly, for each plasmid, two separate tubes were prepared. In one tube, 0.3 µL of Lipofectamine 3000 was diluted and well mixed into 10 µL of Opti-MEM I reduced serum medium (Gibco). In the other tube, 10 µL of Opti-MEM I was first mixed with 0.45 µL of the P3000 reagent by vertexing and then mixed with 45 ng of plasmid DNA (up to 1.75 µL volume) by flicking. The two tubes were then combined, mixed by a brief vertex, and incubated at room temperature for 10 min. For each plasmid, 5 µL of the above DNA/Lipofectamine complex was added to each well for a total of four wells. The amount of plasmid DNA was equivalent to 10 ng for each well, which was determined to be optimal in a separate titration experiment (Fig.2B-C). Then 100 µL of cells (20,000 cells) were added to each well, mixed with the transfection complex, and returned to a CO2 incubator for 48 hours. To measure promoter activity, cells were processed using Promega’s Dual-Luciferase Reporter Assay System. To minimize assay background, all steps were conducted in dark. Firefly luciferase and Renilla luciferase signals were sequentially measured on a GloMax Multi Detection System (Promega). Signal integration time was set to one second per well. Mock transfected cells and empty wells were included to evaluate the assay background.

### Data analysis and statistics

The raw luminescence readouts were processed in Excel in a stepwise manner. First, the Fluc signal was normalized to the corresponding Rluc signal for each well. Second, the average Fluc/Rluc ratio for the no-promoter vector, pLK037, was calculated from its four replicate wells. Third, the Fluc/Rluc ratio of each well was divided by the average pLK037 ratio from step 2 above. This step effectively sets the average Fluc/Rluc ratio of pLK037 to 1, which represents the assay background. Lastly, the normalized promoter activity for each promoter construct was calculated as the average of the normalized Fluc/Rluc ratios among the four replicate wells. The corresponding standard error was calculated as the standard deviation divided by the square root of the number of replicates. Statistical comparison between any two promoter constructs was performed in RStudio using the Basic Statistics and Data Analysis (BSDA) package version 1.2.1, using Welch modified two-sample unpaired t-test assuming unequal variance. Simple linear regression was conducted with the “stats” base package of R version 3.6. The significance level was set at 0.05 for all statistical tests.

## List of abbreviations

5’UTR: 5’ untranslated region
GFP: green fluorescent protein
L1: long interspersed element type 1
M1: monomer 1
M2: monomer 2
M3: monomer 3
MYA: million years ago
non-LTR: non-long terminal repeat.

## Supplemental Information

**Additional file 1:** Promoter constructs and corresponding sequences. Table S1, Promoters assayed in Figure 2. Table S2, Promoters assayed in Figure 3. Table S3, Promoters assayed in Figure 4. Table S4, Table S4. Copy number of two-monomer promoters and individual monomer and tether domains in the mouse genome.

**Additional file 2:** Sequence alignments and cell line authentication report. Figure S1, Alignment of M2 from A_I, Gf_I and Tf_I subfamilies. Figure S2, Alignment of A_I monomers. Figure S3, Alignment of Tf_I monomers. Figure S4, Alignment of Gf_I monomers. Figure S5, Alignment of tether sequences. Figure S6, Cell line authentication report for NIH/3T3.

## Acknowledgements

We thank Arin Smit and Stephan Boissinot for providing mouse L1 consensus sequences, and all An lab members for their enduring support throughout this project.

## Authors’ contributions

LK, KS, JT, CH and CY performed experiments; YH, LW, XG and PY conducted computational analyses; LK and WA designed the project and wrote the manuscript; SN and WA directed the project.

## Availability of data and materials

All data generated or analyzed during this study are available from the corresponding author on reasonable request.

## Competing interests

The authors declare that they have no competing interests.

## Funding

The work was supported by National Institutes of Health [grant numbers R15GM131263 and R03HD099412]. W.A. was supported, in part, by South Dakota State University Markl Faculty Scholar Fund.

## Notes

### Competing Interest Statement

The authors have declared no competing interest.

