## Additional file 2 for "Subfamily-specific differential contribution of individual monomers and the tether sequence to mouse L1 promoter activity"

#### Monomer 2 alignments

```
                 10        20        30        40        50
A_I_M2          GTGCCTGCCCCAATCCAATCGCGCGGAACT-TGAGACTGCGG
Tf_I_M2  GACAGCCGGCCACCTTCCGGACCAGAGGACAGGTGCCCGCCCGGCTGGGG
Gf_I_M2                       TGAGAGCACGGGGGTCTGCCTGGCCTGAG

                 60        70        80        90       100
A_I_M2   TA----CATAGGGAAGCAGGCTACC-CGGGCCTGATCTGGGGCACAAGTC
Tf_I_M2  AGGCGGCCTAAGCCA-CAG-CAGCAGCGGTCGCCATCTTGG-TCCGGGAC
Gf_I_M2  AG---GTTTGTGGCA-CAGGCGCCGGCGGGAGCCTTCTTGGCTCCGGGAC

                110       120       130       140       150
A_I_M2   CCTTCCGCTC---GACTCGAGACTCGAGCCCCGGGCTACCTTGCCAGCAG
Tf_I_M2  CCGCCGAACTTAGGAAATTAGTCT-GAACAGGTGAGAGGGTGCGCCAGAG
Gf_I_M2  TCCGCG-------GAGGGCAGGCT-GCACGGGTGACCGTGTGGAATACAG

                160       170       180       190       200
A_I_M2   AGTCTTGCCCAACACCCGCAAGGGCCCA-----CACGGGACTCCCC--AC
Tf_I_M2  AAC-CTGACAGCTTCTGGAACAGGCAGAAG---CACAGAGGCGCTGAGGC
Gf_I_M2  AGTGCCAGCCGTTTCTGGGACGGGCGAGAGAGTCGCAGAGCTTCTG-GGC

                210       220       230       240
A_I_M2   GGGACCCTAAG-ACCTCTGGTGAGTGGAACACA
Tf_I_M2  AGCACCCTGTGTGGGCCGGG
Gf_I_M2  GGCGCCATCTTCAGCTCCAGACAGCCGGCCACCTTCCTGG
```

**Figure S1.**

#### Alignment of A_I monomers

```
                   10        20        30        40        50
A_Severynse1992    GTGCCTGCCCCAATCCAATCGCACGGAACTTGAGACTGCGGTACATAGGG
A-I_Schichman1993  GTGCCTGCCCCAATCCAATCGCGCGGAACTCGAGACTGCGGTACATAGGG
A1_Zhou2019        GTGCCTGCCCCAATCCAATCGCGCGGAACTTGAGACTGCGGTACATAGGG
A_I_M3             GTGCCTGCCCCAATCCAATCGCGCGGAACTTGAGACTGCGGTACATAGGG
A_I_M2             GTGCCTGCCCCAATCCAATCGCGCGGAACTTGAGACTGCGGTACATAGGG
A_I_M1             GCGCCTACCCCAATCCAATCGCGTGGAACTTGAGACTGCGGTACATAGGG
```

```
                   60        70        80        90        100
A_Severynse1992    AAGCAGGCTACCCGGGCCTGATCTGGGGCACAAGTCCCTTCCGCTCGACT
A-I_Schichman1993  AAGCAGGCTACCCGGGCCTGATCTGGGGCACAAGTCCCTTCCGCTCGACT
A1_Zhou2019        AAGCAGGCTACCCGGGCCTGATCTGGGGCACAAGTCCCTTCCGCTCGACT
A_I_M3             AAGCAGGCTACCCGGGCCTGATCTGGGGCACAAGTCCCTTCCGCTCGACT
A_I_M2             AAGCAGGCTACCCGGGCCTGATCTGGGGCACAAGTCCCTTCCGCTCGACT
A_I_M1             AAGCAGGCTACCCGGGCTTGATCTGGGGCACAAACCCCTTCCACTCCACT
```

```
                   110       120       130       140       150
A_Severynse1992    CGAGACTCGAGCCCCGGGCTACCTTGCCAGCAGAGTCTTGCCCAACACCC
A-I_Schichman1993  CGAGACTCGAGCCCCGGGCTACCTTGCCAGCAGAGTCTTGCCCAACACCC
A1_Zhou2019        CGAGACTCGAGCCCCGGGCTACCTTGCCAGCAGAGTCTTGCCCAACACCC
A_I_M3             CGAGACTCGAGCCCCGGGCTACCTTGCCAGCAGAGTCTTGCCCAACACCC
A_I_M2             CGAGACTCGAGCCCCGGGCTACCTTGCCAGCAGAGTCTTGCCCAACACCC
A_I_M1             CGAG-------CCCCGG-CTACCTTGCCAGCTGAGTC--GCCTGACACCC
```

```
                   160       170       180       190       200
A_Severynse1992    GCAAGGGTCCACACGGGACTCCCCACGGGACCCTAAGACCTCTGGTGAGT
A-I_Schichman1993  GCAAGGGTCCACACGGGACTCCCCACGGGACCCTAAGACCTCTGGTGAGT
A1_Zhou2019        GCAAGGGCCCACACGGGACTCCCCACGGGACCCTAAGACCTCTGGTGAGT
A_I_M3             GCAAGGGTCCACACGGGACTCCCCACGGGACCCTAAGACCTCTGGTGAGT
A_I_M2             GCAAGGGCCCACACGGGACTCCCCACGGGACCCTAAGACCTCTGGTGAGT
A_I_M1             GCAAGGGCCCACACAGGATTCCACACGTGATCCTAAGACCTCTAGTGAGT
```

```
A_Severynse1992    GGATCACA
A-I_Schichman1993  GGATCACA
A1_Zhou2019        GGAACACA
A_I_M3             GGATCACA
A_I_M2             GGAACACA
A_I_M1             GGAACACA
```

**Figure S2.**

#### Alignment of Tf_I monomers

```
                            10        20        30        40        50
Tf_Naas1998         GACAGCCGGCCACCTTCCGGACCAGAGGACAGGTGCCCGCCCGGCTGGGG
Tf_DeBeradinis1999  GACAGCCGGCCACCTTCCGGACCGGAGGACAGGTGCCCGCCCGGCTGGGG
Tf_I_M3             GACAGCCGGCCACCTTCCGGACCGGAGGACAGGTGCCCGCCCGGCTGGGG
Tf_I_M2             GACAGCCGGCCACCTTCCGGACCAGAGGACAGGTGCCCGCCCGGCTGGGG
Tf_I_M1             GACAGCCGGCCACCTTCCGGACCAGAGGACAGGTGCCCGCCCGGCTGGGG

                            60        70        80        90       100
Tf_Naas1998         AGGCGGCCTAAGCCACAGCAGCAGCGGTCGCCATCTTGGTCC-GGGAC-C
Tf_DeBeradinis1999  AGGCGGCCTAAGCCACAGCAGCAGCGGTCGCCATCTTGGTCC-GGGAC-C
Tf_I_M3             AGGCGACCTAAGCCACAGCAGCAGCGGTCGCCATCTTGGTCC-GGGAC-C
Tf_I_M2             AGGCGGCCTAAGCCACAGCAGCAGCGGTCGCCATCTTGGTCC-GGGAC-C
Tf_I_M1             AGGCGACCTAAGCCACAGCAGCAGCGGTCGCCATCTTGGTCCCGGGACTC

                           110       120       130       140       150
Tf_Naas1998         CGCCGAACTTAGGAAATTAGTCTGAACAGGTGAGAGGGTGCGCCAGAGAA
Tf_DeBeradinis1999  CGCCGAACTTAGGAAATTAGTCTGAACAGGTGAGAGGGTGCGCCAGAGAA
Tf_I_M3             CGCCGAACTTAGGAAATTAGTCTGAACAGGTGAGAGGGTGCGCCAGAGAA
Tf_I_M2             CGCCGAACTTAGGAAATTAGTCTGAACAGGTGAGAGGGTGCGCCAGAGAA
Tf_I_M1             CAAGGAACTTAGGAATTTAGTCTGCTTAGGTGAGAGTCTGTACCA-----

                           160       170       180       190       200
Tf_Naas1998         CCTGACAGCTTCTGGAACAGGCGGAAGCACAGAGGCGCTGAGGCAGCACC
Tf_DeBeradinis1999  CCTGACAGCTTCTGGAACAGGCGGAAGCACAGAGGCGCTGAGGCAGCACC
Tf_I_M3             CCTGACAGCTTCTGGAACAGGCGGAAGCACAGAGGCGCTGAGGCAGCACC
Tf_I_M2             CCTGACAGCTTCTGGAACAGGCAGAAGCACAGAGGCGCTGAGGCAGCACC
Tf_I_M1             CCTGGGAACTGCCAAAGCAA-------CACAGTGT--CTGAGAAAGGTCC

                           210
Tf_Naas1998         CTGTGTGGGCCGGG
Tf_DeBeradinis1999  CTGTGTGGGCCGGG
Tf_I_M3             CTGTGTGGGCCGGG
Tf_I_M2             CTGTGTGGGCCGGG
Tf_I_M1             TGTTTTGGGCC
```

**Figure S3.**

#### Alignment of Gf_I monomers

```
                                   10         20         30         40         50
Gf_Goodier2001   TGAGAGCACGGGGGTCTGCCNGGCCTGAGAGGTTTGTGGCACAGGCGCCG
Gf_I_M2          TGAGAGCACGGGGGTCTGCCTGGCCTGAGAGGTTTGTGGCACAGGCGCCG
Gf_I_M1          TGAGAGCACGGGGGTCTGCCTGGCCTGAGAGGTTTGTGGCACAGGCGCCG

                                   60         70         80         90        100
Gf_Goodier2001   GCGGGAGCCTTCTTGGCTCCGGGACTCCGCGGAGGGCAGGCTGCACGGGT
Gf_I_M2          GCGGGAGCCTTCTTGGCTCCGGGACTCCGCGGAGGGCAGGCTGCACGGGT
Gf_I_M1          GCGGGAGCCTTCTTGGCTCCGGGACTCCGCGGAGGGCAGGCTGCACGGGT

                                  110        120        130        140        150
Gf_Goodier2001   GACCGTGTGGAATACAGAGTGCCAGCCGTTTCTGGGACGGGCGAGAGNNN
Gf_I_M2          GACCGTGTGGAATACAGAGTGCCAGCCGTTTCTGGGACGGGCGAGAGAGT
Gf_I_M1          GACCGTGTGGAATACAGAGTGCCAGCCGTTTCTGGGACGGGCGAGAGAGT

                                  160        170        180        190        200
Gf_Goodier2001   CAGAGNNCTTCTGGG-CGGCGCCATCTTCAGCTCCAGACAGCCGGCCACC
Gf_I_M2          CGCAGAGCTTCTGGG-CGGCGCCATCTTCAGCTCCAGACAGCCGGCCACC
Gf_I_M1          CGCAGAGATTCTGGGGCGGCACCATCCTCAGCTCCAGACAACCAGCCACC

Gf_Goodier2001   TTCCTGG
Gf_I_M2          TTCCTGG
Gf_I_M1          TTCCCGG
```

**Figure S4**

#### Alignment of tethers

```
                                                      10        20        30        40        50
A_I_tether
Tf_I_tether
Gf_I_tether   CAAGAGCCACAGAGCTTCTGAGGCTGCGACATCTTCGACTCCAGACAACT
Gf_I_T249

                                                      60        70        80        90       100
A_I_tether    --------------------------------------------------
Tf_I_tether   --------------------------------------------------
Gf_I_tether   GGCCACCTTCCTGGCCAAAGCAACACAGCTTCTGGGAAAGATCCTGTTTT
Gf_I_T249     --------------CCAAAGCAACACAGCTTCTGGGAAAGATCCTGTTTT

                                                     110       120       130       140       150
A_I_tether    ---------ACTTCTGCCAGGAGTCTGGTTCGAACACCAGATATCTGGGT
Tf_I_tether   -----TTCTTCTTCGGCCAGGAGGA-GGTCCAAATACAAGATATCTGCGC
Gf_I_tether   GGGCCTTCACCTTCAGCCAGGAGGA-GGTCCAAACACCAGATAACTGTAC
Gf_I_T249     GGGCCTTCACCTTCAGCCAGGAGGA-GGTCCAAACACCAGATAACTGTAC

                                                     160       170       180       190       200
A_I_tether    ACCTGCCTTGCAAGAAGAGAGCTTGCCTGCAGAGAATACTCTGCCCACTG
Tf_I_tether   ACCTTCCCTGTAAGAG---AGCTTGCCAGCAGAGAGTGCTCTGAGCACTG
Gf_I_tether   ACCTTCCCTGAGAGAGGAGAGCTTGCCTACAGAGACTGCTCTGACCACTG
Gf_I_T249     ACCTTCCCTGAGAGAGGAGAGCTTGCCTACAGAGACTGCTCTGACCACTG

                                                     210       220       230       240       250
A_I_tether    AAACT--AAGGAGAGTGC-TACCCTCCAGGTCTGCTCATAGAGGCTAACA
Tf_I_tether   AAACTCAGAGGAGAGAATCTGTCTCCCAGGTCTGCTGATAGACGGTAACA
Gf_I_tether   AAACTCAGAGAAGAGAGCTTGTCTCCCACGCCTGCTGATAGAGGGTAACA
Gf_I_T249     AAACTCAGAGAAGAGAGCTTGTCTCCCACGCCTGCTGATAGAGGGTAACA

                                                     260       270       280       290       300
A_I_tether    GAGTCACCTGAAGAACAAGCTCTTAACAGTGACAACTAAAACAGCTAGCT
Tf_I_tether   GAATCACCAGAAGAACAATCTCTAAACAGAGTCAACTATAACTACTAACT
Gf_I_tether   AAATCAACAGAGGAACAATCTTTTAACAAAGACAACTATAACAACTAACT
Gf_I_T249     AAATCAACAGAGGAACAATCTTTTAACAAAGACAACTATAACAACTAACT

                                                     310
A_I_tether    TCAGAGATTACCAG
Tf_I_tether   CCAGAGATTACCAG
Gf_I_tether   CCAGAGATTGCCAG
Gf_I_T249     CCAGAGATTGCCAG
```

**Figure S5**

### Cell Line Authentication Service
#### Mouse STR Profile Report

**FTA Barcode:** MUSA1026

**ATCC Sales Order:** SO0814026

| Test Results for Submitted Sample | | | | | ATCC Reference Database Profile | | | |
|---|---|---|---|---|---|---|---|---|
| **Locus** | **Query Profile: MUSA1026** | | | | **Database Profile: CRL-1658 NIH/3T3; Embryonic Fibroblast; Mouse** | | | |
| **18-3** | 17 | 19 | | | 17 | 19 | | |
| **4-2** | 19.3 | 20.3 | | | 19.3 | 20.3 | | |
| **6-7** | 12 | | | | 12 | | | |
| **19-2** | 11 | 12 | | | 11 | 12 | | |
| **1-2** | 13 | 17 | | | 13 | 17 | | |
| **7-1** | 29 | | | | 29 | | | |
| **1-1** | 10 | | | | 10 | | | |
| **3-2** | 14 | 15 | | | 14 | 15 | | |
| **8-1** | 15 | | | | 15 | | | |
| **2-1** | 9 | | | | 9 | | | |
| **15-3** | 20.3 | | | | 20.3 | | | |
| **6-4** | 15.3 | 16.3 | | | 15.3 | | | |
| **11-2** | 15 | 17 | | | 15 | 17 | | |
| **17-2** | 13 | 14 | | | 13 | 14 | | |
| **12-1** | 20 | | | | 20 | | | |
| **5-5** | 14 | 15 | | | 14 | 15 | | |
| **X-1** | 25 | | | | 25 | | | |
| **13-1** | 16.2 | | | | 16.2 | | | |
| Number of shared alleles between query sample and database profile: | | | | | | | | 26 |
| Total number of alleles in the query sample profile: | | | | | | | | 27 |
| Total number of alleles in the database profile: | | | | | | | | 26 |
| Percent match between the submitted sample and the database profile: | | | | | | | | 98 |

**Explanation of Test Results**
Cell lines with ≥80% match are considered to be related; i.e., derived from a common ancestry. Cell lines with a percent match between a 55 - 80% require further investigation for authentication of relatedness.

☐ The submitted sample profile is an exact match for the following ATCC cell line(s) in the ATCC mouse STR database:

☐ The submitted sample profile is mouse, however a matching reference profile has not previously been established in the ATCC mouse STR database.

☑ The submitted profile is similar to the following ATCC cell line(s): CRL-1658

☐ An STR profile could not be generated from the submitted sample.

**Human and/or African Green Monkey Species Detection**

☐ Human and/or African green monkey has been detected in the submitted sample profile (see attached electropherogram at Human D8 & D4 loci).

**Additional Comments:**

The submitted sample MUSA1026 (NIH/3T3) is similar to ATCC cell line CRL-1658 (NIH/3T3).

**Figure S6**
